# WHAT IS NORMAL? MULTIMODAL CHARACTERIZATION OF NON-DISEASED PEDIATRIC DUODENAL BIOPSIES USING MACHINE LEARNING IMAGE ANALYSIS AND TRANSCRIPTOMICS

**DOI:** 10.64898/2026.09.16.752095

**Authors:** Anum Chotani, Jiebei Liu, Nazanin Moradinasab, Shahzaib Khan, Julia Sessions, S. Fisher Rhoads, Srivastava M. Sanjana, Fatima Zulqarnain, Varun Jain, Shyam Raghavan, Christopher Moskaluk, Adam Greene, Chelsea Marie, Donald Brown, Sana Syed

## Abstract

**Objectives:** Pediatric endoscopy is performed only when clinically indicated, limiting access to healthy duodenal tissue. Biopsies with duodenal no pathologic abnormality (NPA) are often used as controls despite the presence of symptoms or inflammatory disease found elsewhere in the gastrointestinal (GI) tract. We characterized pediatric duodenal NPA tissue across clinical, histologic, cellular, and transcriptomic domains.

**Methods:** Archival duodenal NPA biopsies were obtained with clinical metadata and hematoxylin-and-eosin whole-slide images (WSIs). Duodenal mRNA-seq data were analyzed from a subset of patients with duodenal NPA. Clinical metadata and WSIs underwent machine-learning analysis, cell populations were quantified from WSIs, and RNA-seq data underwent differential expression and pathway-enrichment analyses.

**Results:** The primary cohort included 195 patients with duodenal NPA. Comparisons between patients with non-duodenal GI disease and those with no GI disease showed differences in inflammatory biomarkers and follow-up utilization. Unsupervised clinical clustering identified three clusters with partial enrichment for IBD with colonic inflammation and Eosinophilic Esophagitis (EoE) with esophageal inflammation. Supervised clinical classification showed modest discrimination. WSI clustering showed limited disease-status discrimination, and cell quantification showed no significant group differences. In the separate RNA-seq cohort of 43 patients, differential-expression and pathway-enrichment analyses identified transcriptional and pathway-level differences between disease-status groups.

**Conclusions:** This multi-level characterization indicates that pediatric duodenal NPA tissue should not be treated as a uniform control category. Clinical metadata and transcriptomics revealed clinical and molecular heterogeneity, while histologic and cell analyses showed limited disease-status separation, supporting a refined definition of control tissue.

**Study Highlights:** *What is Known:*

⍰ Pediatric duodenal control tissue is difficult to define because asymptomatic children rarely undergo endoscopy.
⍰ Biopsies with duodenal no pathologic abnormality are often used as controls, even when patients have gastrointestinal symptoms or inflammatory disease outside the duodenum.
⍰ Literature suggests there may be variations in control pediatric duodenal tissue.

*What is New:*

⍰ There are variations in inflammatory markers, GI follow-up visits, repeat endoscopies, and gene expression profiles between patients who have no GI disease versus those with non-duodenal GI disease.
⍰ Digital histology, pathologist review, and cell quantification showed limited disease-status separation rather than robust disease-specific histologic differences.

## Introduction

Pediatric patients undergo invasive esophagogastroduodenoscopy (EGD) only when clinically indicated. Therefore, pediatric duodenal tissue used as control is rarely obtained from truly asymptomatic children. In practice, pediatric biopsies with duodenal no pathologic abnormality (NPA) may be used as controls even when patients have gastrointestinal (GI) symptoms or inflammatory disease elsewhere in the GI tract.^1,2^ By contrast, adult control tissue is often obtained from healthy volunteers.^3^ The designation of a sample as “control” is context-dependent; for example, control samples in environmental enteropathy showed regional variation in small intestine inflammation.^4^ Age-related variation in duodenal intraepithelial lymphocyte populations has also been noted.^5^ Specimen acquisition can also influence molecular profiles, as shown by gene-expression differences among histologically “normal” liver biopsies collected by different approaches.^6^ These observations underscore the need to characterize pediatric duodenal NPA tissue before treating it as a uniform disease-study reference.

Digital pathology and machine learning (ML) provide scalable approaches for quantifying histologic patterns in whole-slide images (WSIs).^7^ In gastrointestinal disease studies, ML has been used to classify duodenal biopsies across disease and control categories.^8^ Wei et al. used deep learning to classify celiac disease, nonspecific duodenitis, and normal duodenal tissue^9^, and Syed et al. used ML to classify pediatric celiac disease, environmental enteropathy and normal duodenal tissue.^10^ However, model interpretability and generalizability depend on how control tissue is defined. If duodenal NPA controls are clinically or molecularly heterogeneous, disease classifiers may learn control-selection artifacts or extra-duodenal disease context rather than disease-specific duodenal histopathology.

Here, we characterize pediatric duodenal NPA tissue across clinical, digital histology, cellular, and transcriptomic domains. We examine whether clinical metadata, WSI-based histology features, cell-type composition, and transcriptomic profiles differ by disease status, and how clinical metadata clusters relate to follow-up inflammatory diagnoses or anatomical sites of extra-duodenal inflammation at the index evaluation.

## Methods

### Patient Population and Data Sources

This study was approved by the Institutional Review Board at the University of Virginia (IRB 20866 & 19466). Consent was waived due to the retrospective nature of the study. Patients younger than 18 years at index EGD at the University of Virginia (UVA) were eligible if endoscopic and histopathologic reports documented duodenal no pathologic abnormality (NPA) (Figure 1A; Figure 2). Two analytic sub cohorts were used: Cohort A and Cohort B. Cohort A included pediatric patients with duodenal NPA, clinical metadata, and archival hematoxylin-and-eosin WSIs. Cohort B was a separate pediatric cohort with duodenal NPA and available duodenal mRNA-seq data for transcriptomic analysis. Chart review was performed by a resident physician (JS) and medical student (VJ) from January through September 2022, with double data entry by a research technician (SFR) and VJ. Clinical data were collected from the GI visit before the initial EGD through subsequent GI visits and EGDs up to chart review. Extracted variables included demographics, anthropometrics, symptoms, past and family history, medication use, endoscopic and pathologic findings, biomarkers, follow-up utilization, and follow-up diagnoses. Descriptive statistics for Cohort A are summarized in **Table 1**. For analyses requiring disease-status labels, patients were assigned to no GI disease or Non-duodenal disease based on documented inflammatory disease status. No GI disease was defined as absence of inflammatory GI disease in past medical history, absence of extra-duodenal inflammation at the index evaluation, and no new inflammatory GI diagnosis during follow-up. Non-duodenal disease was defined as inflammatory GI disease or extra-duodenal GI inflammation documented before the index EGD, at the index evaluation, or during follow-up. Inflammatory GI disease included inflammatory bowel disease, eosinophilic esophagitis, and celiac disease.

**Figure 1.**
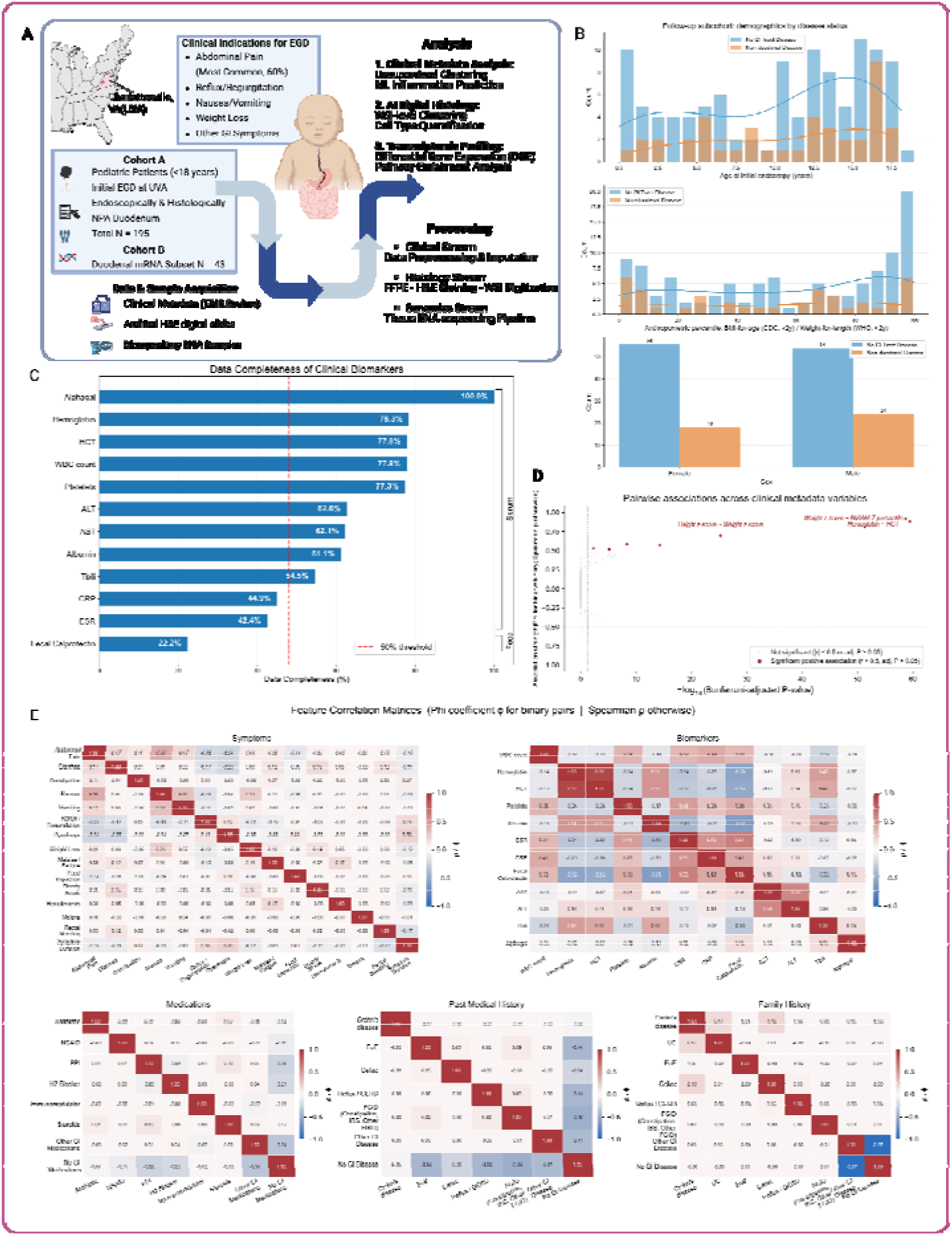
Study workflow and clinical data landscape. (A)Retrospective UVA study design and analytic cohorts. (B) Demographic distributions of the labelable Cohort A subcohort by disease status. (C) Clinical biomarker completeness; fecal calprotectin, ESR, and CRP were flagged with testing-status indicators. (D) Pairwise clinical-variable associations after redundancy filtering. (E) Within-domain correlation matrices for symptoms, biomarkers, medications, past medical history, and family history. NPA, no pathologic abnormality; WSI, whole-slide image.

**Figure 2:**
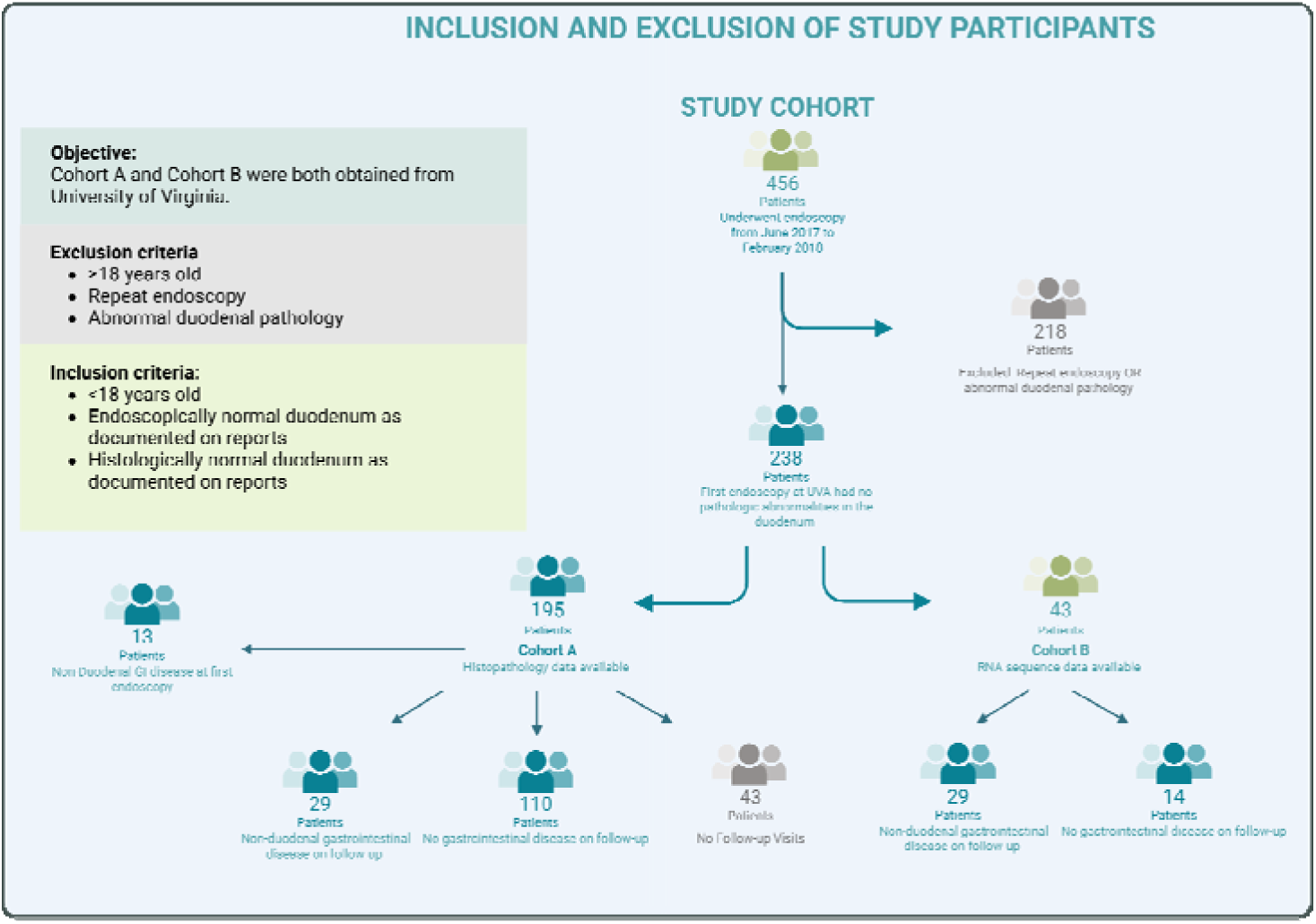
Study Inclusion and Exclusion. Pediatric patients were included if they were less than 18 years of age at the time of an initial EGD at the University of Virginia (UVA) and had an endoscopically and histologically normal duodenum as documented reports. A total of 456 patients were screened, out of these 218 patients either had repeat endoscopy or had abnormal duodenal pathology and these were excluded. The remaining 238 patients were included in the study, out of which 195 patients had digital histopathology data available and 43 patients had RNA sequencing completed.

**Table 1:**
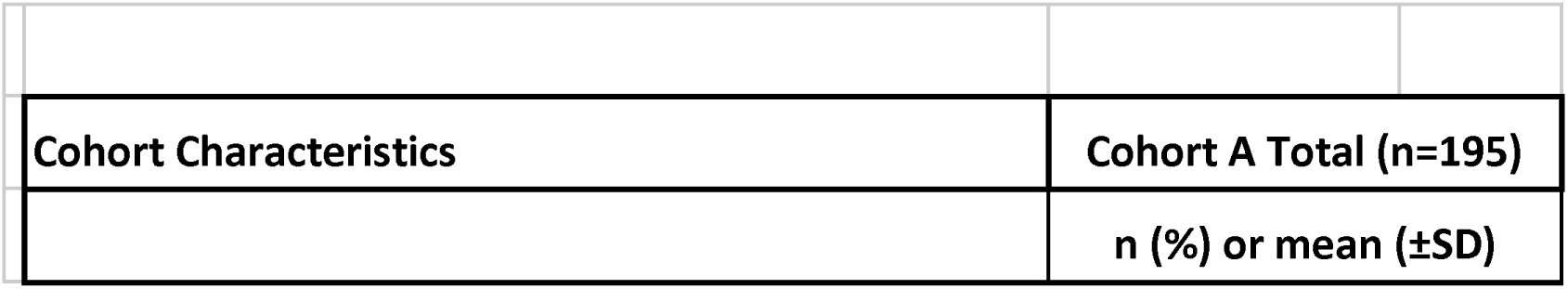
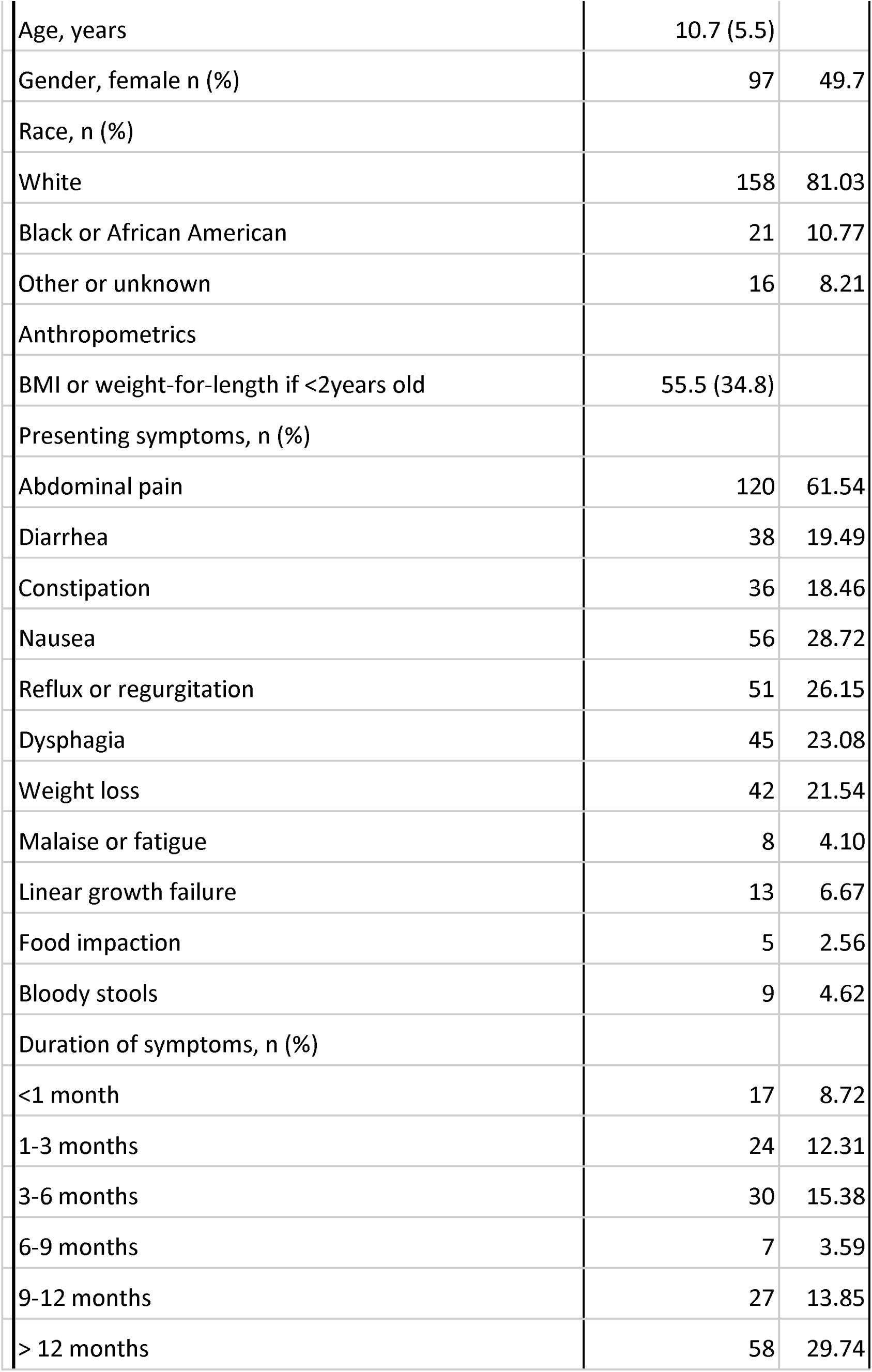
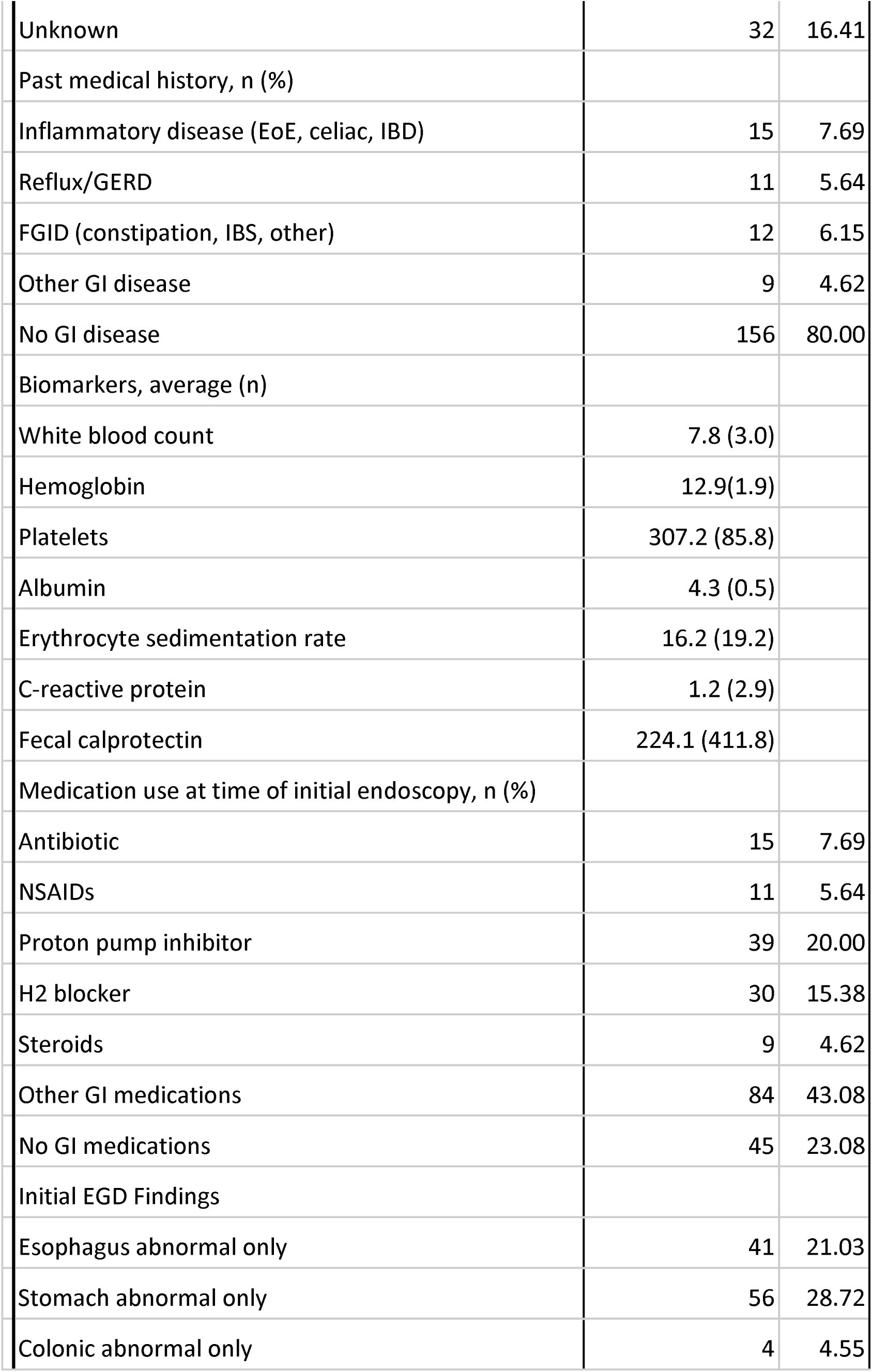
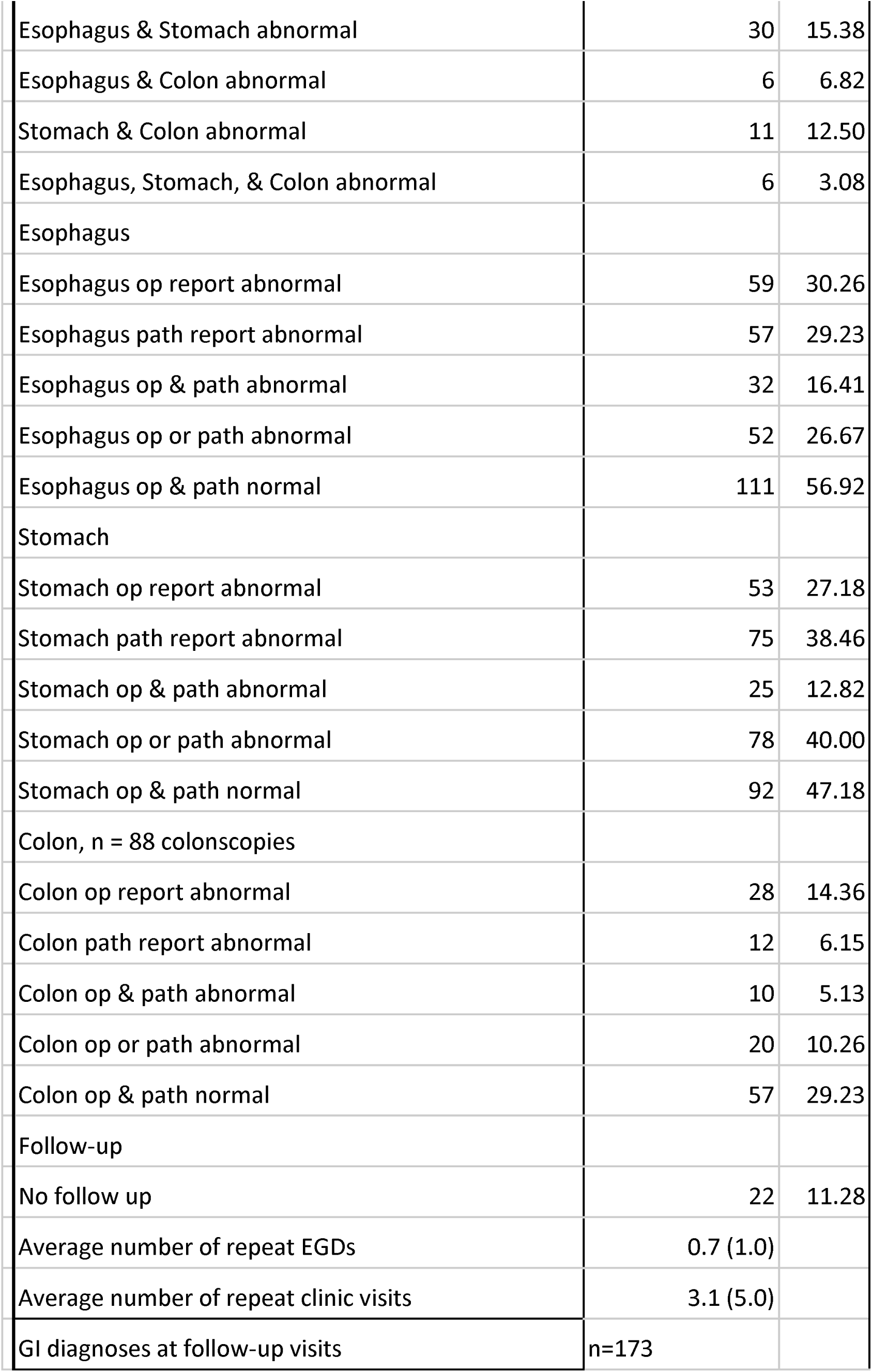
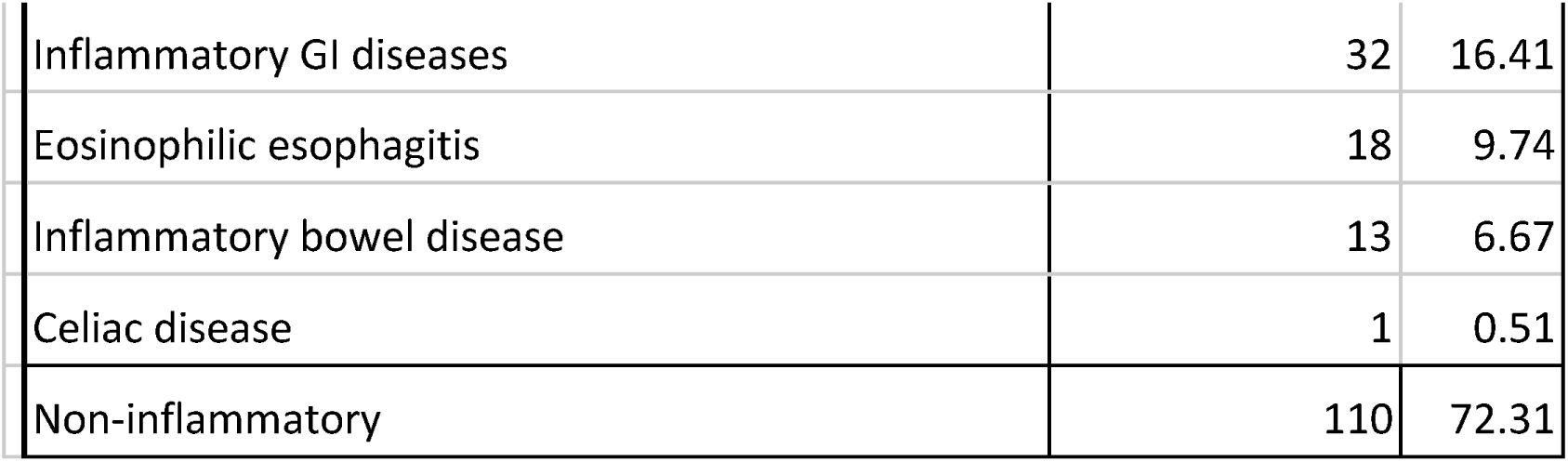
Demographics and clinical metadata for the total cohort for primary cohort A.

|  |  |
| --- | --- |
| Cohort Characteristics | Cohort A Total (n=195) |
|  | n (%) or mean (±SD) |

|  |  |  |
| --- | --- | --- |
| Age, years | 10.7 (5.5) |  |
| Gender, female n (%) | 97 | 49.7 |
| Race, n (%) |  |  |
| White | 158 | 81.03 |
| Black or African American | 21 | 10.77 |
| Other or unknown | 16 | 8.21 |
| Anthropometrics |  |  |
| BMI or weight-for-length if <2years old | 55.5 (34.8) |  |
| Presenting symptoms, n (%) |  |  |
| Abdominal pain | 120 | 61.54 |
| Diarrhea | 38 | 19.49 |
| Constipation | 36 | 18.46 |
| Nausea | 56 | 28.72 |
| Reflux or regurgitation | 51 | 26.15 |
| Dysphagia | 45 | 23.08 |
| Weight loss | 42 | 21.54 |
| Malaise or fatigue | 8 | 4.10 |
| Linear growth failure | 13 | 6.67 |
| Food impaction | 5 | 2.56 |
| Bloody stools | 9 | 4.62 |
| Duration of symptoms, n (%) |  |  |
| <1 month | 17 | 8.72 |
| 1-3 months | 24 | 12.31 |
| 3-6 months | 30 | 15.38 |
| 6-9 months | 7 | 3.59 |
| 9-12 months | 27 | 13.85 |
| > 12 months | 58 | 29.74 |

|  |  |  |
| --- | --- | --- |
| Unknown | 32 | 16.41 |
| Past medical history, n (%) |  |  |
| Inflammatory disease (EoE, celiac, IBD) | 15 | 7.69 |
| Reflux/GERD | 11 | 5.64 |
| FGID (constipation, IBS, other) | 12 | 6.15 |
| Other GI disease | 9 | 4.62 |
| No GI disease | 156 | 80.00 |
| Biomarkers, average (n) |  |  |
| White blood count | 7.8 (3.0) |  |
| Hemoglobin | 12.9(1.9) |  |
| Platelets | 307.2 (85.8) |  |
| Albumin | 4.3 (0.5) |  |
| Erythrocyte sedimentation rate | 16.2 (19.2) |  |
| C-reactive protein | 1.2 (2.9) |  |
| Fecal calprotectin | 224.1 (411.8) |  |
| Medication use at time of initial endoscopy, n (%) |  |  |
| Antibiotic | 15 | 7.69 |
| NSAIDs | 11 | 5.64 |
| Proton pump inhibitor | 39 | 20.00 |
| H2 blocker | 30 | 15.38 |
| Steroids | 9 | 4.62 |
| Other GI medications | 84 | 43.08 |
| No GI medications | 45 | 23.08 |
| Initial EGD Findings |  |  |
| Esophagus abnormal only | 41 | 21.03 |
| Stomach abnormal only | 56 | 28.72 |
| Colonic abnormal only | 4 | 4.55 |

|  |  |  |
| --- | --- | --- |
| Esophagus & Stomach abnormal | 30 | 15.38 |
| Esophagus & Colon abnormal | 6 | 6.82 |
| Stomach & Colon abnormal | 11 | 12.50 |
| Esophagus, Stomach, & Colon abnormal | 6 | 3.08 |
| Esophagus |  |  |
| Esophagus op report abnormal | 59 | 30.26 |
| Esophagus path report abnormal | 57 | 29.23 |
| Esophagus op & path abnormal | 32 | 16.41 |
| Esophagus op or path abnormal | 52 | 26.67 |
| Esophagus op & path normal | 111 | 56.92 |
| Stomach |  |  |
| Stomach op report abnormal | 53 | 27.18 |
| Stomach path report abnormal | 75 | 38.46 |
| Stomach op & path abnormal | 25 | 12.82 |
| Stomach op or path abnormal | 78 | 40.00 |
| Stomach op & path normal | 92 | 47.18 |
| Colon, n = 88 colonoscopies |  |  |
| Colon op report abnormal | 28 | 14.36 |
| Colon path report abnormal | 12 | 6.15 |
| Colon op & path abnormal | 10 | 5.13 |
| Colon op or path abnormal | 20 | 10.26 |
| Colon op & path normal | 57 | 29.23 |
| Follow-up |  |  |
| No follow up | 22 | 11.28 |
| Average number of repeat EGDs | 0.7 (1.0) |  |
| Average number of repeat clinic visits | 3.1 (5.0) |  |
| GI diagnoses at follow-up visits | n=173 |  |

|  |  |  |
| --- | --- | --- |
| Inflammatory GI diseases | 32 | 16.41 |
| Eosinophilic esophagitis | 18 | 9.74 |
| Inflammatory bowel disease | 13 | 6.67 |
| Celiac disease | 1 | 0.51 |
| Non-inflammatory | 110 | 72.31 |

### Clinical Metadata Analysis

Clinical metadata were analyzed using pairwise association analysis, k-means clustering with gastroenterologist review, and TabPFN^11^. Disease-status labels, endoscopic or histopathologic disease-defining variables, and follow-up-utilization variables, including repeat clinic visits and repeat EGDs, were excluded from clustering and classifier inputs.

Data preparation included recoding laboratory values below or above assay thresholds(i.e., CRP <0.1 was recoded to 0.1), excluding sparsely reported variables including ferritin, GGT, and INR, collapsing celiac testing into positive, negative, and not-tested categories, and removing highly correlated or redundant variables (Supplementary Table 1). For fecal calprotectin, ESR, and CRP, which had greater than 50% missingness, binary testing-status indicators were created to encode clinically informative missingness. Pairwise associations were evaluated using Spearman’s ρ for continuous-variable pairs and signed Phi coefficients for binary-variable pairs, with Bonferroni correction. Within-domain correlation matrices were generated for symptoms, biomarkers, medications, past medical history, and family history.

#### Clustering

Unsupervised k-means clustering was performed on standardized clinical metadata from Cohort A. Continuous fecal calprotectin, ESR, and CRP values were excluded from clustering, while their binary testing-status indicators were retained. Missing values in the remaining clustering features were imputed using K-nearest neighbors (KNN) after feature scaling. Candidate k values were assessed using the SSE elbow, silhouette score, Calinski–Harabasz index, and Davies–Bouldin index; the final three-cluster solution was selected based on internal clustering metrics and clinical interpretability. Presenting symptoms were compared across clusters using chi-squared tests with Bonferroni correction. Biomarker differences were assessed using Kruskal–Wallis tests on unscaled observed values with Benjamini–Hochberg correction. Clusters were then assessed for overlap with follow-up diagnoses and anatomical sites of extra-duodenal inflammation.

#### Classification

TabPFN^11^ was performed to distinguish Non-duodenal disease from No GI disease using clinical metadata. The same disease-status, disease-defining, and follow-up-utilization exclusions were applied. Missing values were handled natively by TabPFN. Testing-status indicators for fecal calprotectin, ESR, and CRP were retained. Five-fold stratified cross-validation was performed. Feature attribution was assessed using SHAP analysis^29^.

### Histology Image Analysis

Archival hematoxylin and eosin (H&E)-stained duodenal biopsies from the initial EGD were digitized by the Biorepository and Tissue Research Facility (BTRF) at UVA using a Hamamatsu NanoZoomer S360 Digital slide scanner S1322-01. One biopsy per patient was selected, generating two WSIs. WSIs were partitioned into non-overlapping 512×512-pixel patches using a sliding-window approach. A ResNet-18–based autoencoder^12^ was used to extract patch embeddings (Figure 3B). Patch embeddings from both WSIs were assigned to a patch-level Gaussian Mixture Model (GMM), and cluster-membership probabilities were aggregated into normalized patient-level histograms. These histograms were clustered using a second GMM, with cluster number selected by Bayesian Information Criterion. WSI-derived clusters were reviewed by two gastrointestinal pathologists (SR and CM) to identify cluster-defining histologic, anatomic, or technical features. A Random Forest classifier was trained on patient-level WSI histograms to distinguish Non-duodenal disease from No GI disease and evaluated using five-fold cross-validation.

**Figure 3.**
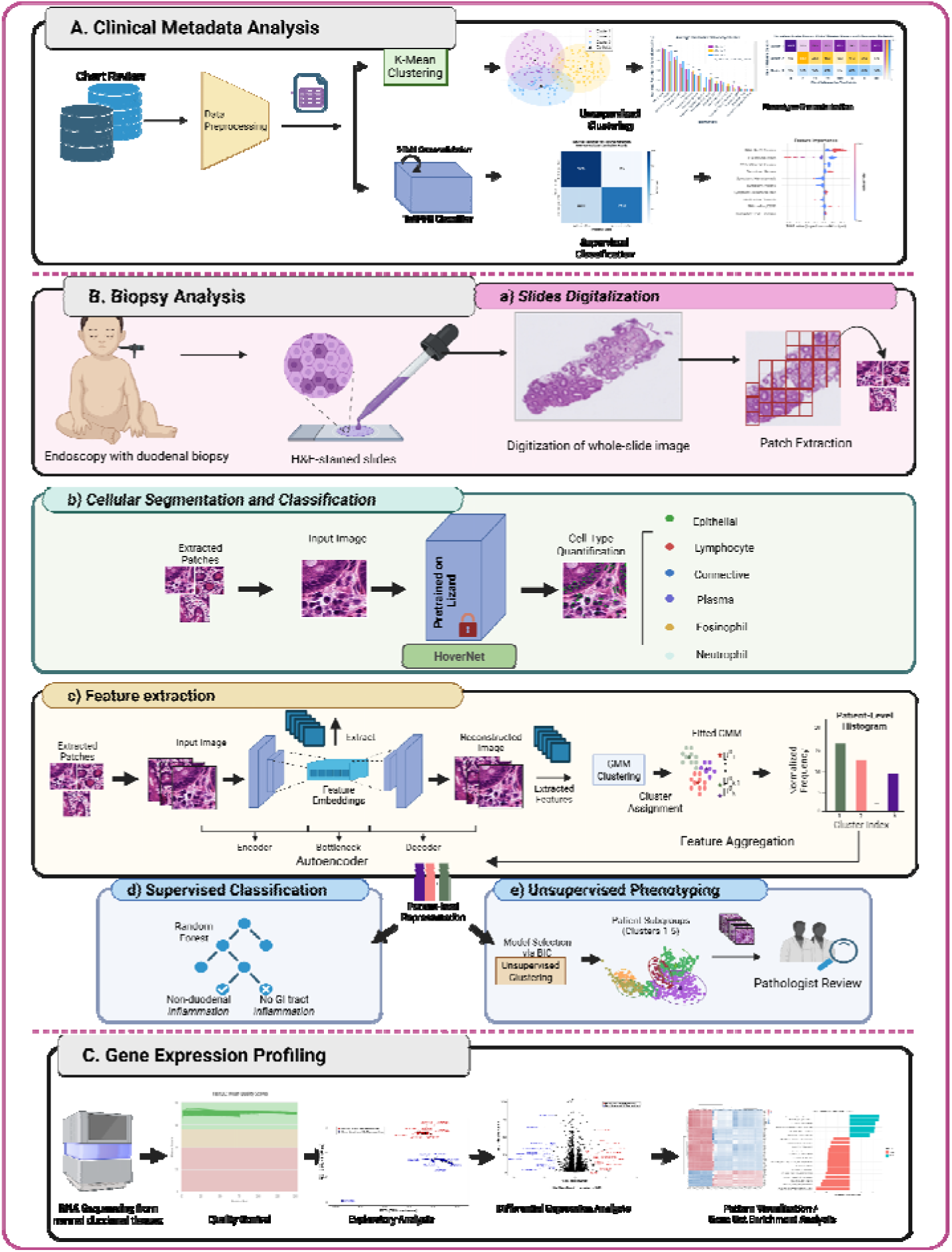
Analytical workflow for multimodal characterization of pediatric duodenal NPA tissue. (A) Clinical metadata underwent preprocessing, feature filtering, pairwise association analysis, k-means clustering, gastroenterologist review, and TabPFN classification; disease-defining and follow-up-utilization variables were excluded from model inputs. (B) H&E duodenal WSIs were tiled for HoVerNet cell quantification and ResNet-18 autoencoder feature extraction. GMM-derived patient histograms were used for Random Forest classification and unsupervised phenotyping with pathologist review. (C) Separate duodenal mRNA-seq data underwent quality control, differential expression, visualization, and GSEA.

#### Cell Type Quantification

Nuclear segmentation and classification were performed using HoVerNet^13^ pretrained on the Lizard dataset^14^. Patch-level predictions from both WSIs were aggregated per patient to quantify epithelial cells, lymphocytes, plasma cells, neutrophils, eosinophils, and connective-tissue cells (fibroblasts, muscle, and endothelial cells). Cell populations were summarized as patient-level relative abundance and density.

### Gene Expression Profiling

A separate pediatric duodenal NPA cohort with mRNA-seq data was analyzed. This dataset was originally collated as a control group for a study of environmental enteropathy.^15^ mRNA-seq bioinformatics analysis was performed by the UVA Bioinformatics Core, RRID:SCR_012718. The workflow is summarized in Figure 3C, with detailed methods in Supplementary Information A. RNA-seq analyses included quality control, mapping, quantitation, batch correction, differential gene-expression, and pathway-enrichment analysis. Differential expression was evaluated between Non-duodenal disease and No GI disease. Normalized expression data were used for principal-component analysis and heatmap visualization. Pathway enrichment was assessed using Gene Set Enrichment Analysis against MSigDB Hallmark gene sets, with DESeq2 Wald statistic gene ranking and Benjamini–Hochberg-adjusted P values.

## Results

### Patient Characteristics

A total of 456 pediatric patients underwent EGD at UVA between July 2017 and February 2018. After excluding 218 patients with repeat EGD or abnormal duodenal endoscopic or histopathologic findings, 238 patients with duodenal NPA were eligible for study inclusion (Figure 2). Cohort A included 195 patients with clinical metadata and archival H&E WSIs. Cohort B included 43 separate patients with duodenal mRNA-seq data for transcriptomic analysis. In Cohort A, the mean age was 10.7 years; 49.7% of patients were female, and 81.0% were White. Demographic, biomarker, diagnostic, endoscopic, and follow-up data are summarized in Table 1. Within Cohort A, 152 patients had sufficient follow-up or documentation for disease-status classification: 110 were classified as No GI disease and 42 as Non-duodenal GI disease, including 13 with prior non-duodenal GI disease at the index EGD and 29 diagnosed with inflammatory disease during follow-up. The most common presenting symptom was abdominal pain (60%); the most frequent symptom-duration category was greater than 12 months (29%). Initial endoscopy findings for Cohort A showed that 57% had a normal-appearing esophagus and 47% had a normal-appearing stomach. Among the 90 patients who underwent same-day colonoscopy, 63% had a normal-appearing colon and ileum.

### Clinical Metadata Analysis

Clinical biomarker completeness varied; fecal calprotectin, ESR, and CRP fell below 50% completeness and were represented by testing-status indicators for downstream modeling (**Figure 1c**). Pairwise association analysis after redundancy filtering showed structured relationships among clinical variables (**Figure 1d**). Within-domain correlation matrices further summarized associations within symptoms, biomarkers, medications, past medical history, and family history (**Figure 1e**).

#### Clustering

Unsupervised k-means clustering of Cohort A clinical metadata identified three clinical clusters: Cluster 1 (n=69), Cluster 2 (n=83), and Cluster 3 (n=43) (**Figure 4a**). Presenting symptoms and observed biomarker profiles differed across clusters (**Figures 4b** and **4c**). Demographic characteristics are summarized in **Supplementary Table 2**. Follow-up diagnoses showed partial cluster enrichment: IBD cases were concentrated in Cluster 1 (12/14, 86%), EoE cases in Cluster 2 (14/19, 74%), and the single celiac disease case also fell in Cluster 2 (**Figure 4d**). In the column-normalized anatomical-site analysis, all colonic-only cases and most colon-involved mixed-site cases mapped to Cluster 1 (C, 100%; SC, 82%; ESC, 83%). Esophageal-only cases were enriched in Cluster 2 (E, 66%; **Figure 4e**).

**Figure 4.**
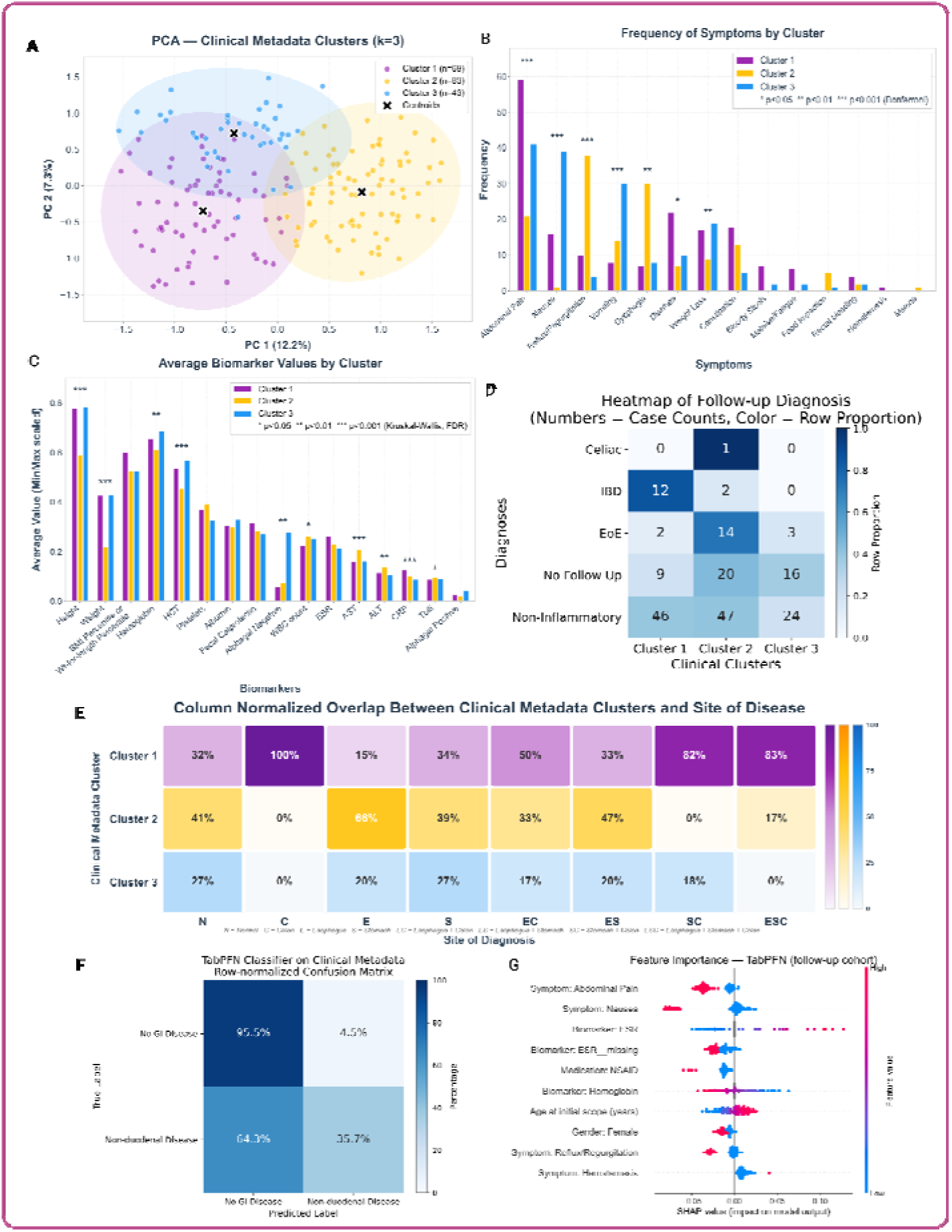
Clinical metadata clustering and classification. (A)PCA visualization of k-means clusters from a reduced clinical-feature set excluding disease-defining, follow-up-utilization, and continuous high-missingness biomarker variables. (B) Presenting symptoms by cluster. (C) Biomarker profiles by cluster. (D) Follow-up diagnoses across clusters. (E) Column-normalized concordance between clusters and anatomical sites of extra-duodenal inflammation. (F) Row-normalized TabPFN confusion matrix for Non-duodenal disease versus No GI disease. (G) SHAP feature-attribution summary. Asterisks indicate corrected P values as described in Methods.

#### Classification

TabPFN classification modestly distinguished No GI disease from Non-duodenal disease, with 78.9% accuracy, an F1 score of 0.484, high specificity, and limited sensitivity (TN=105, FP=5, FN=27, TP=15; **Figure 4f**). SHAP analysis identified abdominal pain, nausea, and inflammatory biomarker-related features as major contributors to TabPFN predictions (**Figure 4g**). These features should be interpreted as exploratory model-attribution signals rather than independent clinical predictors.

### Histology Image Analysis

Digitization of one selected biopsy per Cohort A patient generated 390 WSIs, with two WSIs per patient. WSIs were partitioned into non-overlapping 512 × 512-pixel patches, yielding 197,944 patches. Unsupervised clustering of patient-level histograms identified five histology clusters (n=5, 88, 32, 3, and 67; **Supp Figure 1a**). Histology clusters did not show a clear pattern by anatomical site of macroscopic or microscopic extra-duodenal inflammation at the index evaluation (**Supp Figure 1b**). Follow-up diagnoses showed no clear enrichment across histology clusters, although EoE cases were most frequently assigned to Cluster 2 (52.7%; **Supp Figure 1d**). Pathologist review showed that cluster-defining features were predominantly technical or anatomical rather than disease-specific: Cluster 1 showed detached epithelial regions consistent with tissue-handling artifact; Cluster 2 showed villi with tangentially cut crypts, consistent with sectioning orientation; Cluster 3 showed long, slender finger-like villi; Cluster 4 showed detached villous fragments consistent with tissue-handling artifact; and Cluster 5 showed mucosa with prominent Brunner glands, consistent with duodenal bulb sampling (**Supp Figure 1e**).

#### Classification

A Random Forest classifier trained on patient-level WSI histograms modestly distinguished Non-duodenal disease from No GI disease in the labelable subcohort. Using five-fold cross-validation, the model achieved 67.8% accuracy and an F1 score of 0.33 (TN=91, FP=19, FN=30, TP=12; Supp Figure 1c).

#### Cell Type Quantification

**Supplementary Figure 2** shows WSI-derived patient-level relative abundance and density of epithelial cells, lymphocytes, plasma cells, neutrophils, eosinophils, and connective-tissue cells. No cell type differed significantly between No GI disease and Non-duodenal disease groups after Bonferroni correction. Cell-type distributions were also stable across anatomical sites of macroscopic or microscopic inflammation (**Supplementary Figure 3**). Cell-type summaries also did not differ significantly across clinical metadata clusters (**Supplementary Figure 4**). In contrast, cell-type composition differed across histology image clusters, consistent with the technical, anatomical, and tissue-structure differences identified by pathologist review rather than disease-status separation (**Supplementary Figure 5**).

### Gene Expression Profiling

Cohort B included 43 pediatric patients with duodenal NPA and duodenal mRNA-seq data, including 29 with Non-duodenal disease and 14 with No GI disease;demographic characteristics are summarized in **Supplementary Table 3**. PCA of normalized expression data showed separation between Non-duodenal disease and No GI disease samples along PC1; PC1 and PC2 explained 75% and 24% of variance, respectively (**Supp Figure 6**). Differential-expression analysis identified genes upregulated in No GI disease, including IFI6, IFIT1, MSMB, IFI44L and C3, and in Non-duodenal disease, including PDE4C, SLC11A2 and IGHG4 (**Supp Figure 6**). The top 100 differentially expressed genes showed group-associated expression patterns by hierarchical clustering (**Supp Figure 6**). Hallmark pathway analysis showed enrichment of interferon-α/γ response and proliferation-related pathways in No GI disease, and oxidative phosphorylation, fatty-acid metabolism, bile-acid metabolism, and adipogenesis in Non-duodenal disease (**Supp Figure 6**).

## Discussion

This study characterizes pediatric duodenal no pathologic abnormality (NPA) tissue across clinical metadata, digital histology, cell-type quantification, and transcriptomic profiling. This work aligns with Human Cell Atlas and the Chan Zuckerberg Initiative pediatric tissue-mapping efforts to define cellular and molecular reference states.^16–18^ As the proximal small intestine and a common biopsy site in pediatric gastrointestinal evaluation, the duodenum is an important context for defining appropriate controls. Our findings indicate that pediatric duodenal NPA tissue should not be treated as a uniform control category.

Clinical metadata demonstrated heterogeneity among patients with pediatric duodenal NPA. Abdominal pain and prolonged symptom duration were common before the index evaluation. Unsupervised clustering identified three clinical metadata clusters with distinct symptom and biomarker profiles. Cluster 1 was enriched for IBD-associated follow-up diagnoses and colonic inflammation, whereas Cluster 2 was enriched for EoE-associated diagnoses and esophageal inflammation. Symptoms and biomarker categories varied across clusters, including gastrointestinal symptoms, anthropometrics, hematological markers, and inflammatory biomarkers (**Figure 2b** and **2c**). Because follow-up-utilization variables were excluded from clustering inputs, these associations are less likely to reflect clustering by subsequent care intensity alone. However, the overlap between clinical clusters and disease status was incomplete, indicating that clinical metadata captured broad phenotypic structure rather than discrete diagnostic categories.

Supervised classification using clinical metadata modestly distinguished No GI disease from Non-duodenal disease. The model showed high specificity but limited sensitivity, indicating that clinical metadata alone were insufficient to identify all patients with extra-duodenal inflammatory disease. SHAP analysis highlighted symptoms and inflammatory biomarker-related features as contributors to model predictions; however, these findings should be interpreted as exploratory attribution signals rather than independent clinical predictors. These results support the value of careful pre-procedural assessment while emphasizing that clinical metadata cannot replace endoscopic or histologic evaluation.

Digital histology showed limited disease-status separation in duodenal H&E WSIs. Although unsupervised image clustering identified five patient-level histology clusters, pathologist review indicated that cluster-defining features were predominantly technical or anatomical, including tissue-handling artifact, sectioning orientation, villous morphology, and duodenal bulb sampling. Random Forest classification using patient-level WSI histograms also showed limited disease-status discrimination. These findings suggest that duodenal NPA tissue from patients with and without extra-duodenal disease may appear similar by routine H&E morphology, and that image-based models may learn preparation- or sampling-related features if control definitions and slide-level quality factors are not carefully specified.

Cell-type quantification further supported limited histologic separation between disease-status groups. HoVerNet-derived cell summaries did not differ significantly between No GI disease and Non-duodenal disease after multiple-comparison correction. Cell-type distributions were also stable across anatomical sites of extra-duodenal inflammation. In contrast, cell composition differed across histology image clusters, consistent with the technical, anatomical, and tissue-structure differences rather than disease-status separation.

Prior machine-learning studies have also reported difficulty distinguishing control or histologically normal duodenal tissue from disease. In our group’s previous work, misclassifications were most frequent between celiac disease and normal duodenal tissue.^19^ Similarly, Khan et al. reported limited accuracy for classifying normal tissue against celiac disease or environmental enteropathy.^20^ Our pediatric duodenal NPA WSIs with nuclear annotations provide a detailed reference for future model development. However, such models should account for patient-level clinical context, slide-preparation artifacts, and anatomic sampling variation. Control tissue in gastrointestinal pathology should therefore be defined by both duodenal histopathology and broader clinical disease status.

Transcriptomic analysis identified molecular differences in a separate pediatric duodenal NPA cohort. Although all samples had duodenal NPA, normalized mRNA-seq profiles separated No GI disease from Non-duodenal disease along the first principal component. Differential-expression and pathway-enrichment analyses showed group-associated transcriptional patterns, with interferon-response and proliferation-related pathways enriched in No GI disease and metabolic pathways in Non-duodenal disease. These findings indicate that tissue classified as duodenal NPA by routine histopathology may still show molecular heterogeneity associated with disease context.

Future studies should validate these findings in larger, multi-center pediatric cohorts with harmonized clinical metadata, prospective follow-up, matched histology and transcriptomic data, and more diverse disease groups. Such studies should evaluate whether refined control-tissue definitions improve model robustness and interpretability. Less invasive approaches for characterizing intestinal biology may also broaden access to pediatric reference cohorts beyond children undergoing clinically indicated endoscopy. Future work should explicitly define controls by both tissue-level histopathology and patient-level clinical context.

This study has several limitations. First, analyses were retrospective and single-center, and the clinical, WSI, and RNA-seq data were not all obtained from the same patients. Second, disease-status classification depended on available follow-up or documented inflammatory disease status, which may introduce ascertainment bias. Third, the RNA-seq cohort had limited sex representation, limiting generalizability of transcriptomic findings. Fourth, disease-status groups were imbalanced, particularly in supervised classification, and model-attribution results should be interpreted cautiously. Finally, duodenal WSIs contained preparation and sampling variation, including tissue fragmentation, sectioning-orientation differences, and duodenal bulb versus distal duodenal sampling, which influenced unsupervised image clusters.

As machine-learning models are increasingly developed for clinical pathology and gastroenterology, attention to bias, reproducibility, and external validation is essential. Single-center pediatric models may reflect local referral patterns, documentation practices, tissue processing, and sampling variation rather than generalizable disease biology. Future model-development studies should include multi-center validation, transparent preprocessing and modeling workflows, and, where feasible, shared code, model documentation, and de-identified derived data. These safeguards are particularly important when using control tissues that are histologically NPA but clinically heterogeneous.

## Acknowledgements

The authors thank the UVA Bioinformatics Core (RRID:SCR_012718) for their assistance with the gene expression data.

## Ethics

This study was approved by the Institutional Review Board at the University of Virginia (IRB 20866 & 19466). Consent was waived due to the retrospective nature of the study.

## Abbreviations

ALT: Alanine Aminotransferase
BTRF: Biorepository and Tissue Research Facility
CRP: C-Reactive Protein
DGE: Differential Gene Expression
EGD: Esophagogastroduodenoscopy
EoE: Eosinophilic Esophagitis
ESR: Erythrocyte Sedimentation Rate
GI: Gastrointestinal
GMM: Gaussian Mixture Model
H&E: Hematoxylin and Eosin
IBD: Inflammatory Bowel Disease
INR: International Normalized Ratio
IRB: Institutional Review Board
KNN: K-Nearest Neighbors
ML: Machine Learning
mRNA-seq: Messenger RNA Sequencing
MSigDB: Molecular Signatures Database
NPA: No Pathologic Abnormality
PCA: Principal Component Analysis
RNA-seq: RNA Sequencing
SHAP: SHapley Additive exPlanations
SSE: Sum of Squared Errors
UVA: University of Virginia
WBC: White Blood Cell
WSI: Whole-Slide Image

## Supplementary Information

### A. Detailed Methods Used for Gene Expression Profiling

#### Quality Control

Samples had an average of approximately 25 million paired-end reads, sufficient for gene-level quantification. Read quality was assessed using FastQC program (http://www.bioinformatics.babraham.ac.uk/projects/fastqc/) and raw data quality report was generated using the MultiQC tool.^21^

#### Mapping and Quantitation

Reads were aligned using the splice-aware aligner STAR^22^. Prior to mapping, a human reference index was constructed based on the GRCh38 human genome reference (GRCh38.d1.vd1.fa & gencode.v27.annotation.gtf) with the sjdboverhang parameter set to 149 to match the read length of our samples. More than 95% of reads mapped to the human reference genome. Gene-based read counts were derived from the aligned reads and a count matrix was generated, which served as the input file for the analysis of differential gene expression.

#### Batch Correction and Differential Gene Expression Analysis (DGE)

The DESeq2 package was used to conduct differential gene expression analysis.^23^ Lowly expressed genes were filtered out. Following data normalization, PCA plots were generated to observe the unsupervised clustering of control and disease samples. Normalized data (scaled to sequencing depth) was used for batch correction using the sva for sequencing (svaseq) batch correction tool.^24^ Surrogate variable decomposition was calculated for the data and used with the sample condition for modeling. Surrogate variable analysis was performed using the sva package to account for unwanted variation while preserving the disease-status contrast. The number of surrogate variables was estimated using the Leek method. Identified surrogate variables were included as covariates in the DESeq2 design model for differential expression analysis. Principal-component analysis was performed on normalized transformed expression values to visualize separation between No GI disease and Non-duodenal disease samples. Lowly expressed genes were excluded from the analysis before identifying differential gene expression. Data normalization, dispersion estimates, and model fitting (negative binomial) were carried out with the DESeq function. The log-transformed, normalized gene expression of the top 500 most variable genes was used to perform an unsupervised principal component analysis. Differentially expressed genes were summarized using log2 fold change estimates and Benjamini–Hochberg false-discovery-rate adjusted P values. Log2 fold-change estimates were shrunken using the apeglm method through the DESeq2 lfcShrink function^25^. The MA plot and Volcano plot represent significantly up- and down-regulated genes, and were generated using the specific functions in the DESeq2 package. The heat map was generated using the pheatmap package in R.

#### Pathway Analysis

Pathway analysis was performed using the fgsea package in Bioconductor R (https://bioconductor.org/packages/release/bioc/html/fgsea.html). The reference database for pathway enrichment analysis comprised Hallmark, C2, and C5 gene sets from the msigdb.^26–28^ Pathways were ranked by normalized enrichment score, and statistical significance was assessed using adjusted P values. Additionally, GSEA-style plots were generated for both up- and down-regulated pathways.

**Supp Figure 1.**
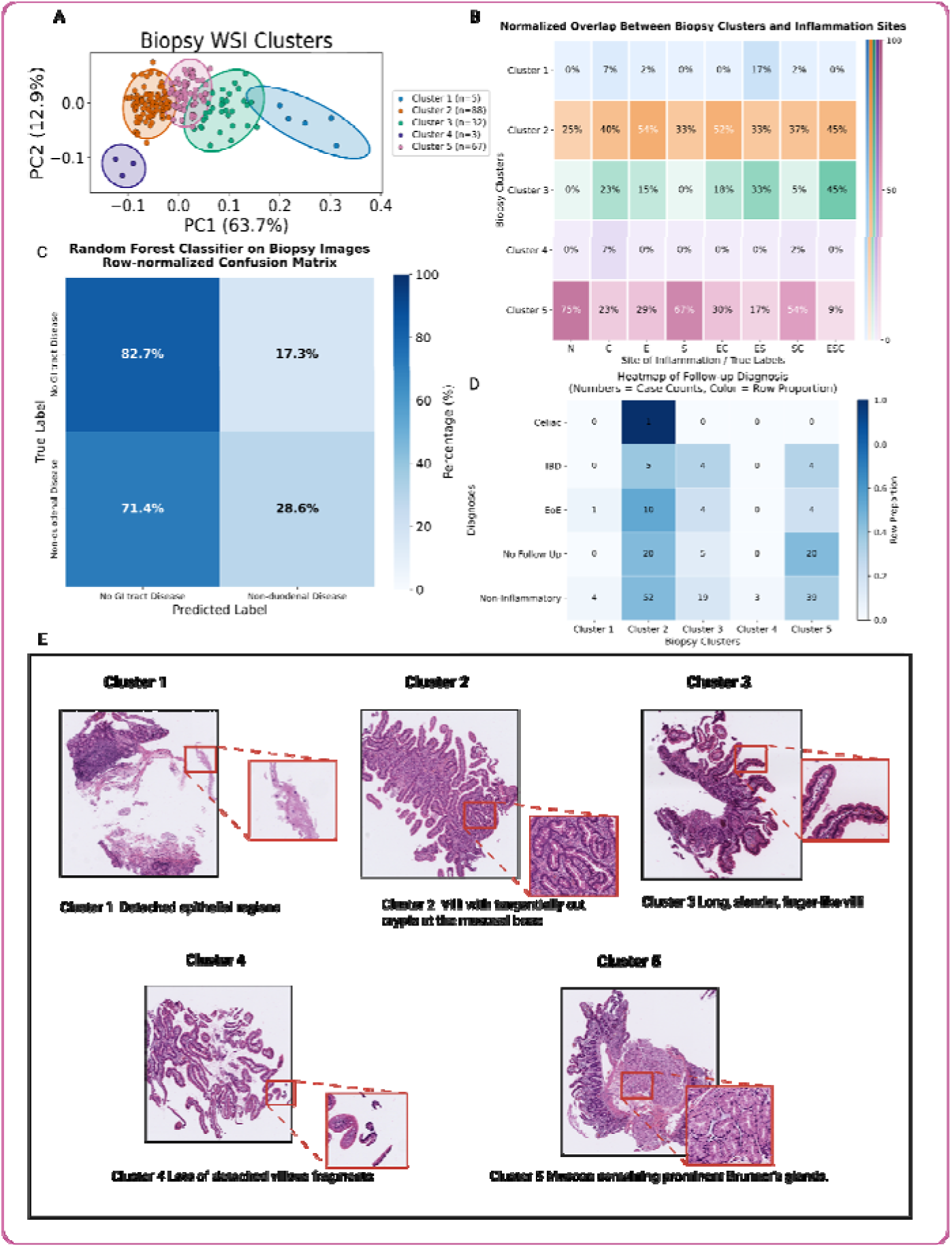
Duodenal histology clustering and classification. (A)Patient-level Gaussian Mixture Model clustering of WSI patch-embedding histograms from Cohort A (n=195; k=5 by BIC). (B) Concordance between histology clusters and anatomical sites of extra-duodenal inflammation. (C) Random Forest classification of Non-duodenal disease versus No GI disease in the labelable subcohort (n=152). (D) Follow-up diagnoses by histology cluster. (E) Representative H&E patches; cluster-defining features were predominantly technical or anatomical rather than disease-specific histopathologic findings. WSI, whole-slide image; BIC, Bayesian Information Criterion.

**Supplementary Figure 2.**
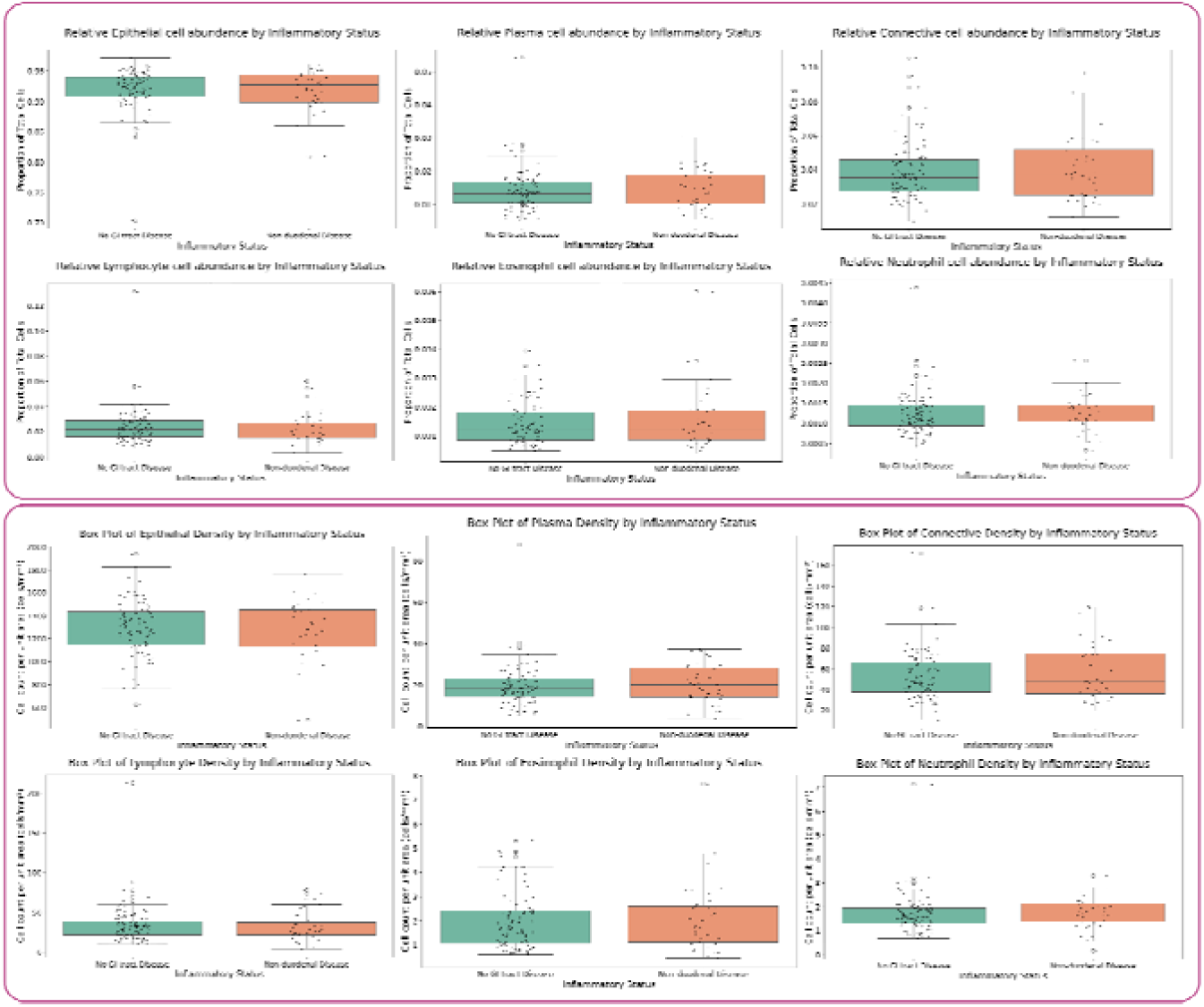
Cell-population quantification by disease-status group. Boxplots show WSI-derived patient-level relative abundance and density of six HoVerNet-identified cell types in the labelable subcohort: No GI disease (n=110) and Non-duodenal disease (n=42). Cell types included epithelial cells, lymphocytes, plasma cells, neutrophils, eosinophils, and connective-tissue cells. Boxes indicate median and interquartile range; whiskers indicate 1.5 × IQR. Group comparisons used Wilcoxon rank-sum tests with Bonferroni correction across 12 tests. No significant disease-status differences were detected.

**Supplementary Figure 3.**
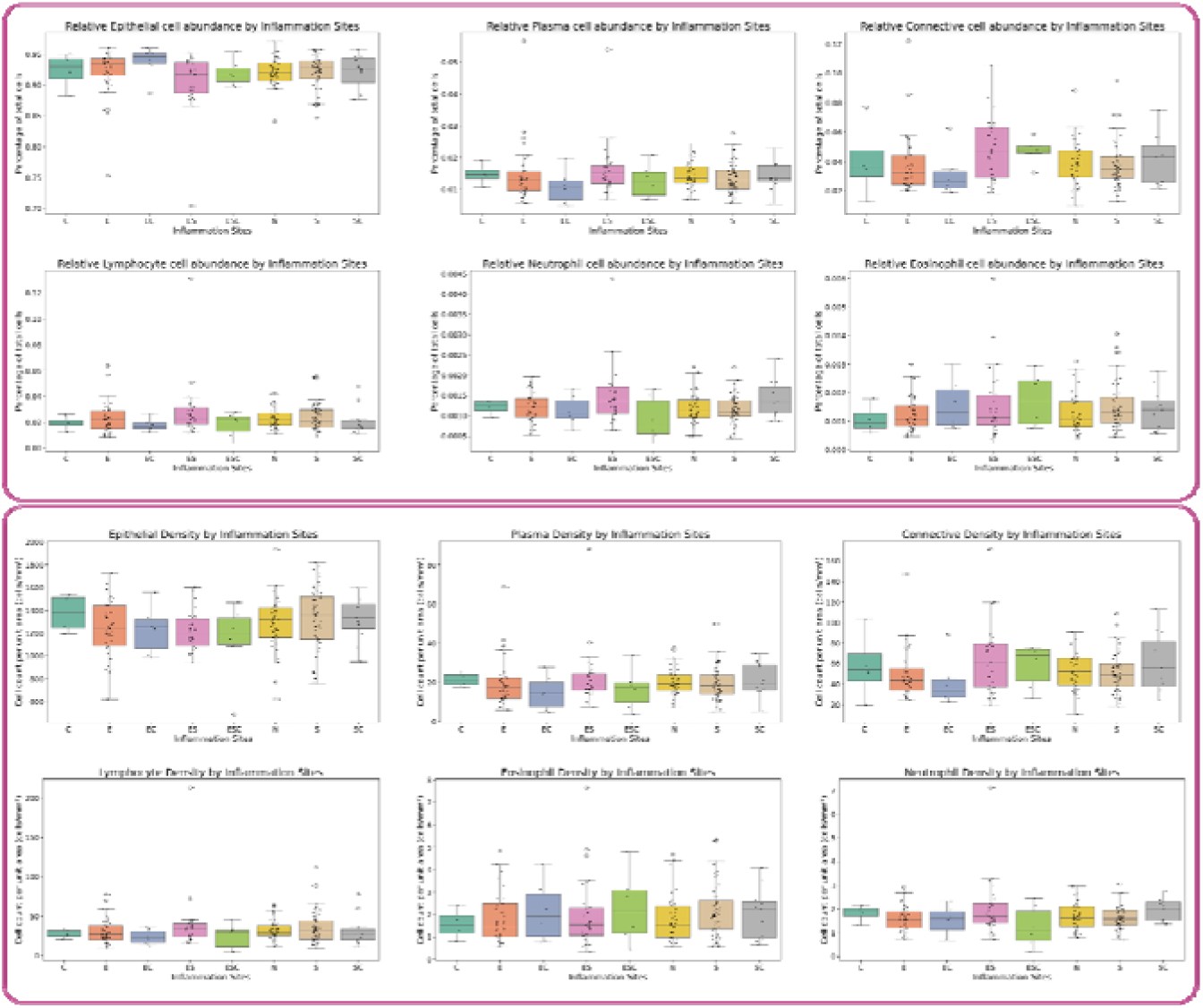
Cell-type composition by anatomical site of extra-duodenal inflammation. Boxplots show WSI-derived patient-level relative abundance and density of HoVerNet-identified cell types across anatomical sites of macroscopic or microscopic inflammation. No significant cell-type differences were detected after multiple-comparison correction. Site labels indicate location of inflammation: C, colon; E, esophagus; EC, esophagus and colon; ES, esophagus and stomach; ESC, esophagus, stomach, and colon; N, no pathologic abnormality; S, stomach; SC, stomach and colon.

**Supplementary Figure 4.**
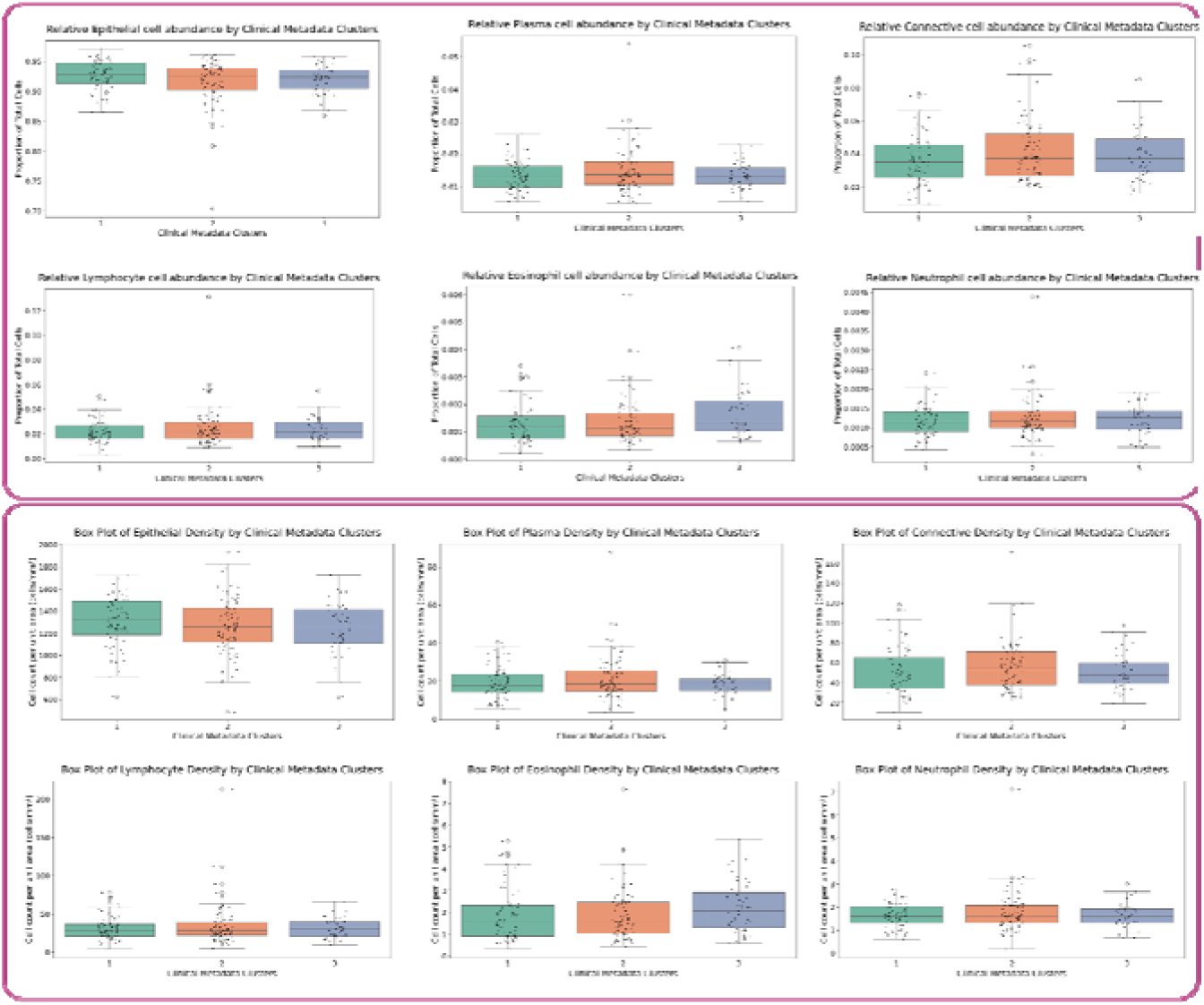
Cell-type composition across clinical metadata clusters. Boxplots show WSI-derived patient-level relative abundance and density of HoVerNet-identified epithelial cells, lymphocytes, plasma cells, neutrophils, eosinophils, and connective-tissue cells across the three k-means clinical metadata clusters. Statistical comparisons used Kruskal–Wallis tests. No significant cell-type differences were detected across clinical metadata clusters, indicating limited correspondence between unsupervised clinical phenotypes and duodenal cellular composition.

**Supplementary Figure 5.**
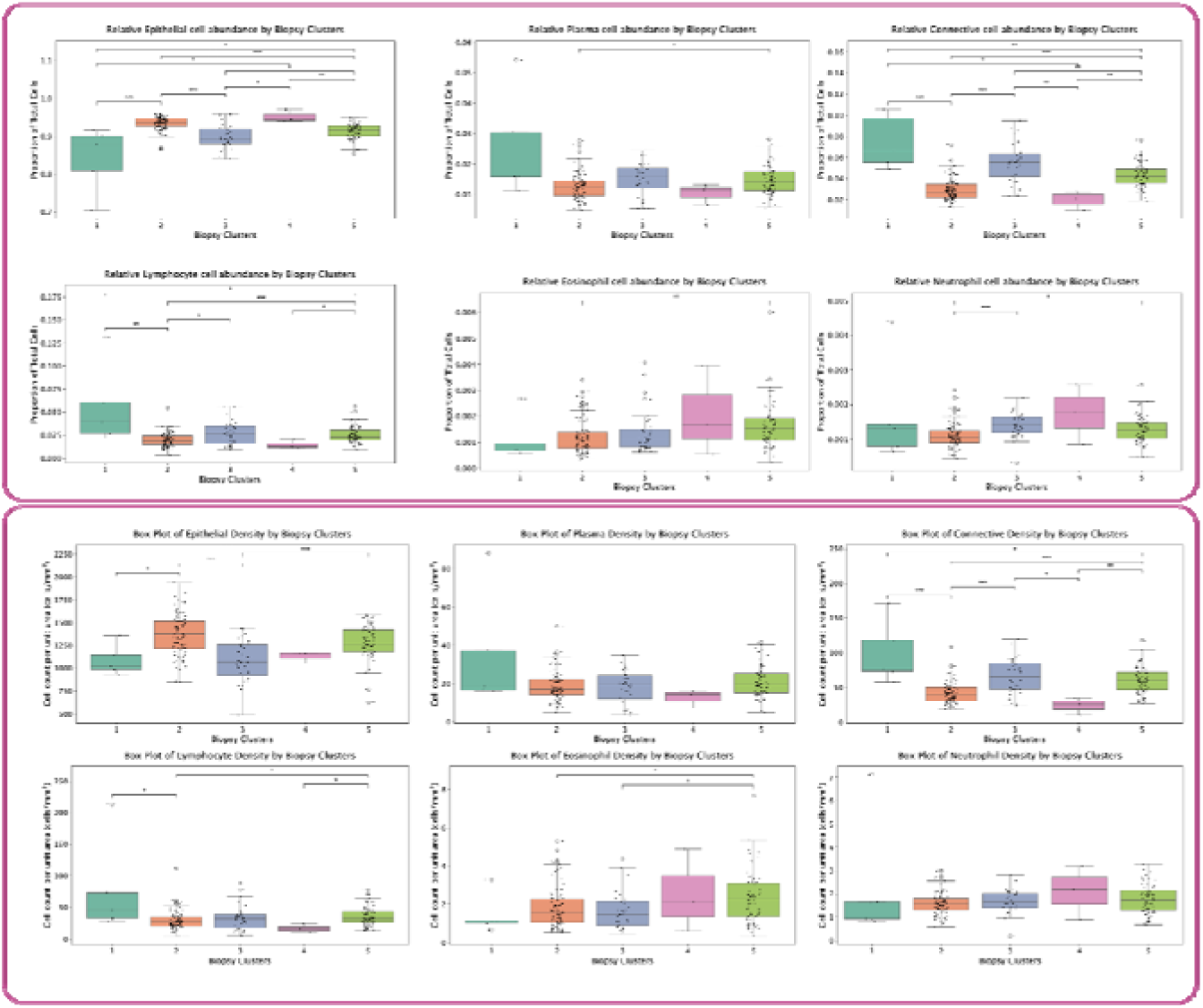
Cell-type composition across histology image clusters. Boxplots show WSI-derived patient-level relative abundance and density of HoVerNet-identified cell types across the five unsupervised histology image clusters. Cell-type composition differed significantly across image clusters, consistent with tissue-structure, anatomic-sampling, and slide-preparation variation identified by pathologist review rather than disease-status separation. Asterisks denote statistical significance: *P < 0.05, **P < 0.01, ***P < 0.001.

**Supp Figure 6.**
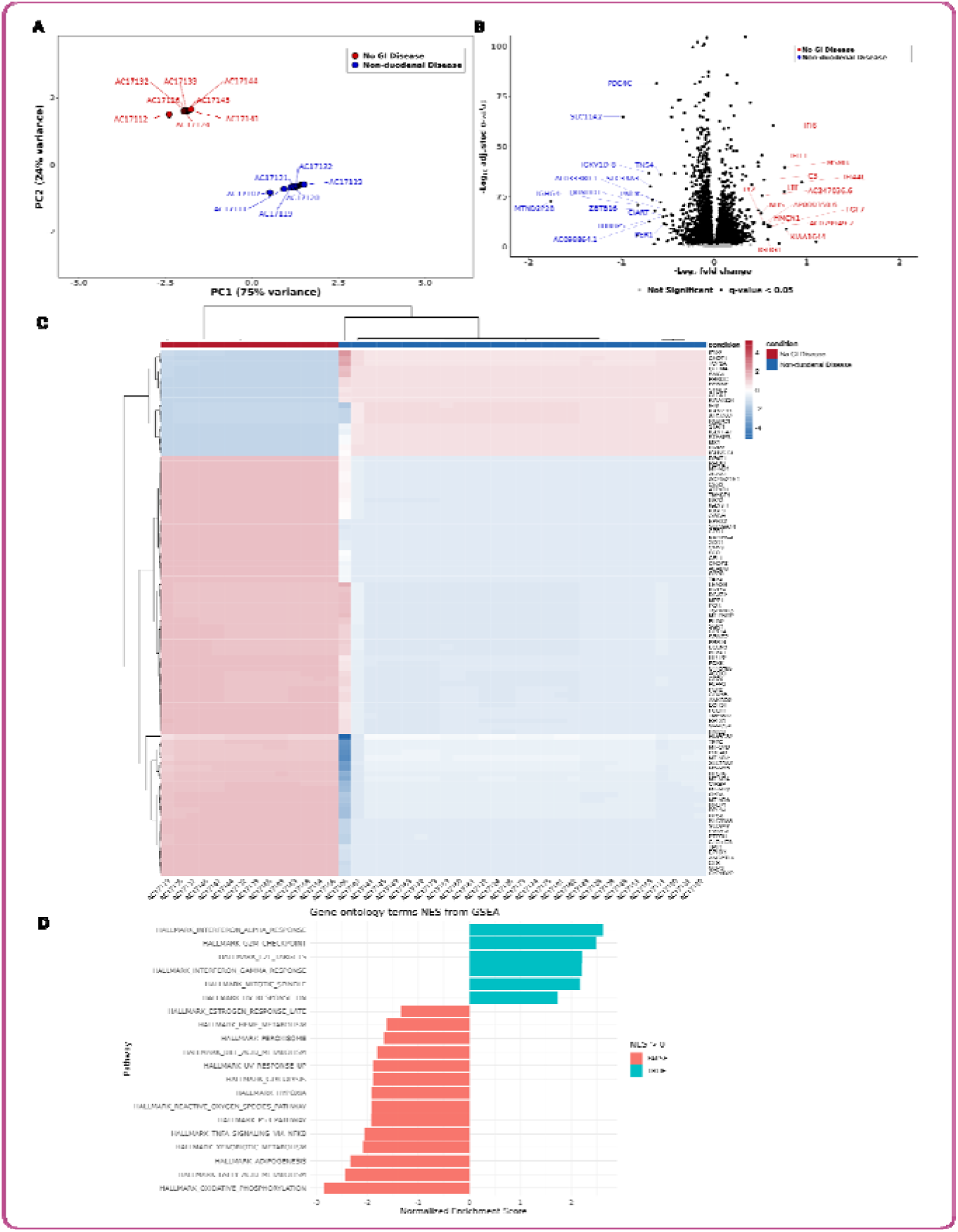
Gene-expression profiling of pediatric duodenal NPA tissue.(less than 100 words) (A)PCA of normalized mRNA-seq profiles from Cohort B, comparing No GI disease and Non-duodenal disease groups. (B) Volcano plot showing differentially expressed genes; red labels indicate genes upregulated in No GI disease, and blue labels indicate genes upregulated in Non-duodenal disease. (C) Heatmap of the top 100 differentially expressed genes. (D) Hallmark pathway enrichment ranked by normalized enrichment score. NPA, no pathologic abnormality; NES, normalized enrichment score.

**Supplemental Table 1.** Table of clinical metadata used in unsupervised hierarchical clustering and clinical classification. Abbreviations: Erythrocyte Sedimentation Rate (ESR), C-Reactive Protein (CRP), Aspartate Transferase (AST), Alanine Aminotransferase (ALT), Proton Pump Inhibitor (PPI), gastroesophageal reflux disease (GERD), Functional Gastrointestinal Disorder (FGID), Non-Steroidal Anti-Inflammatory Drugs (NSAIDs). * = variables removed due to redundancy, high correlation, or non-informative/all-zero encoding. ** = variables imputed by indicator above 50% missingness. † Follow-up-utilization variables excluded from clustering and classifier inputs.

| Demographics | Anthropometrics | Past Medical History (PMH) | Family History | Symptoms | Biomarkers | Medications |
| --- | --- | --- | --- | --- | --- | --- |
| Age at initial scope (years) | Height* | Crohn's disease | Crohn's disease | Abdominal pain | WBC | Antibiotics |
| Gender | Height Percentile* | Ulcerative colitis* | Ulcerative colitis | Diarrhea | Hemoglobin | NSAIDs |
| Race | Height z-score* | Unspecified IBD* | Unspecified IBD | Constipation | Hematocrit | PPI |
| Ethnicity | Weight* | Eosinophilic esophagitis | Eosinophilic esophagitis | Nausea | Platelets | H2 Blockers |
|  | Weight Percentile* | Celiac disease | Celiac disease | Vomiting | Albumin | Immunomodulators |
|  | Weight z-score* | Inflammatory diseases* | Inflammatory diseases* | Reflux/Regurgitation | ESR** | Steroids |
|  | BMI* | Reflux/GERD | Reflux/GERD | Dysphagia | CRP** | Other GI meds |
|  | BMI Percentile* | FGID (Constipation, IBD, Other FGID) | FGID (Constipation, IBD, Other FGID) | Weight loss | Fecal calprotectin** | No GI meds |
|  | BMI z-score* | Other GI disease | Other GI disease | Malaise/fatigue | AST |  |
|  | BMI Percentile or Wt-for-length Percentile | No GI disease | No GI disease | Linear growth failure | ALT |  |
|  |  |  |  | Food impaction | Total bilirubin |  |
|  |  |  |  | Bloody stools | Alpha gal positive |  |
|  |  |  |  | Hematemesis | Alpha gal negative |  |
|  |  |  |  | Melena |  |  |
|  |  |  |  | Rectal bleeding |  |  |
|  |  |  |  | Symptom duration |  |  |
|  |  |  |  | # of Repeat EGDs† |  |  |
|  |  |  |  | # of Repeat Clinic Visits† |  |  |

**Supplemental Table 2:**
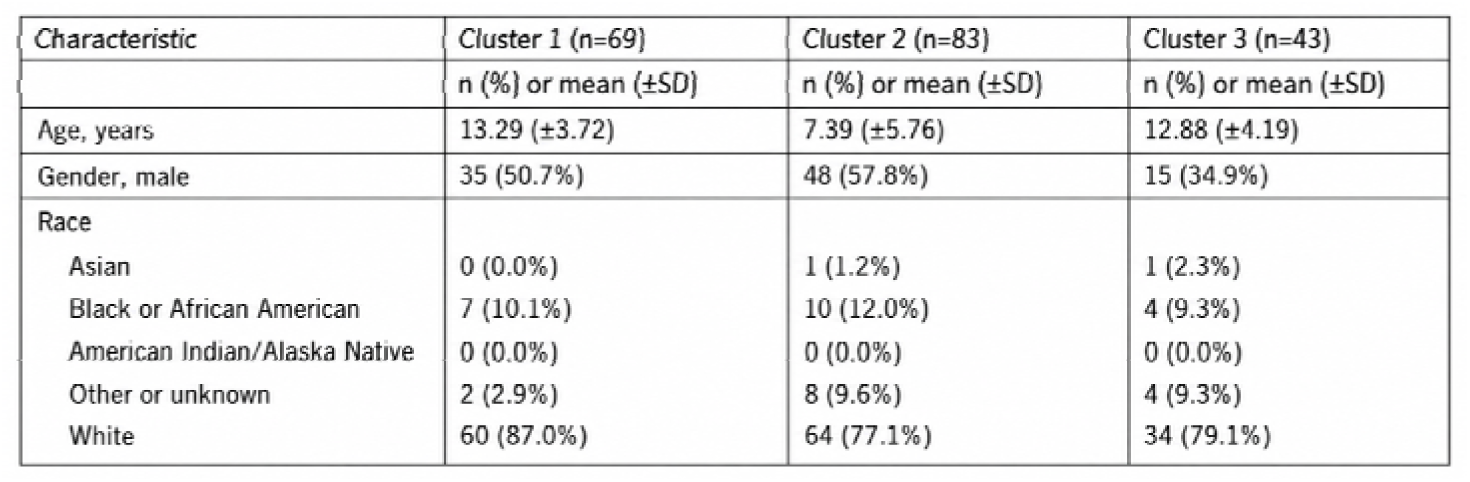
Demographic data for each of the 3 unsupervised clinical metadata clusters.

**Supplemental Table 3:**
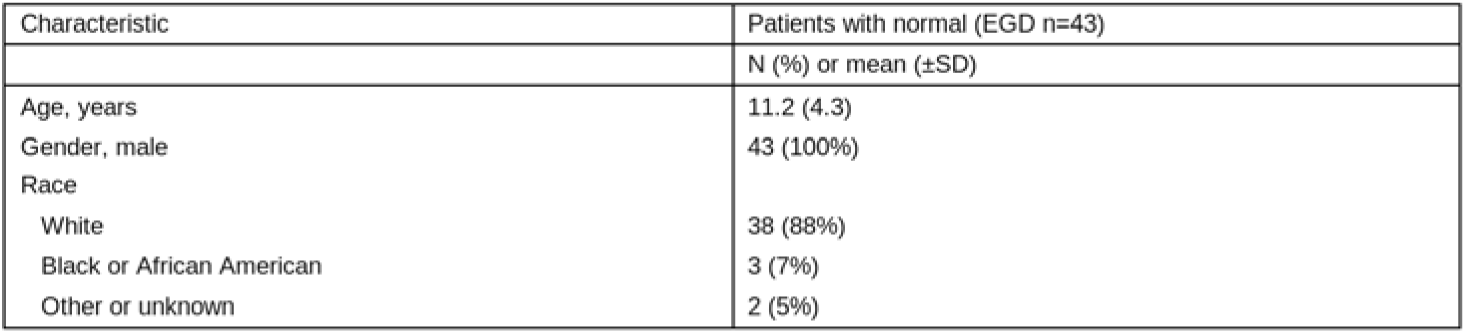
Demographic data for the secondary cohort B used for gene expression profiling.

| Characteristic | Patients with normal (EGD n=43) |
| --- | --- |
| | N (%) or mean (±SD) |
| Age, years | 11.2 (4.3) |
| Gender, male | 43 (100%) |
| Race |  |
| White | 38 (88%) |
| Black or African American | 3 (7%) |
| Other or unknown | 2 (5%) |

## References

1. Quinn L, Nguyen B, Menard-Katcher C, Spencer L. IgG4+ cells are increased in the gastrointestinal tissue of pediatric patients with active eosinophilic gastritis and duodenitis and decrease in remission. Digestive and Liver Disease. 2023;55(1):53–60. doi:10.1016/j.dld.2022.08.020

2. Loberman-Nachum N, Sosnovski K, Di Segni A, et al. Defining the Celiac Disease Transcriptome using Clinical Pathology Specimens Reveals Biologic Pathways and Supports Diagnosis. Sci Rep. 2019;9(1):1–10. doi:10.1038/s41598-019-52733-1

3. Bruce JK, Burns GL, Sinn Soh W, et al. Defects in NLRP6, autophagy and goblet cell homeostasis are associated with reduced duodenal CRH receptor 2 expression in patients with functional dyspepsia. Brain Behav Immun. 2022;101(December 2021):335–345. doi:10.1016/j.bbi.2022.01.019

4. Watanabe K, Petri WA. Environmental Enteropathy: Elusive but Significant Subclinical Abnormalities in Developing Countries. EBioMedicine. 2016;10:25–32. doi:10.1016/j.ebiom.2016.07.030

5. Camarero C, Leon F, Sanchez L, Asensio A, Roy G. Age-related variation of intraepithelial lymphocytes subsets in normal human duodenal mucosa. Dig Dis Sci. 2007;52(3):685–691. doi:10.1007/s10620-006-9176-3

6. Asselah T, Bièche I, Laurendeau I, et al. Significant gene expression differences in histologically Normal liver biopsies: Implications for control tissue. Hepatology. 2008;48(3):953–962. doi:10.1002/hep.22411

7. Lecun Y, Bengio Y, Hinton G. Deep learning. Nature. 2015;521(7553):436–444. doi:10.1038/nature14539

8. Popovic D, Glisic T, Milosavljevic T, et al. The Importance of Artificial Intelligence in Upper Gastrointestinal Endoscopy. Diagnostics. 2023;13(18):1–18. doi:10.3390/diagnostics13182862

9. Wei JW, Wei JW, Jackson CR, Ren B, Suriawinata AA, Hassanpour S. Automated Detection of Celiac Disease on Duodenal Biopsy Slides: A Deep Learning Approach. J Pathol Inform. 2019;9(1):1–7. doi:10.4103/jpi.jpi_87_18

10. Syed S, Al-Boni M, Khan MN, et al. Assessment of Machine Learning Detection of Environmental Enteropathy and Celiac Disease in Children. JAMA Netw Open. 2019;2(6):e195822. doi:10.1001/jamanetworkopen.2019.5822

11. Hollmann, Noah, et al. “Tabpfn: A transformer that solves small tabular classification problems in a second.” arXiv preprint arXiv:2207.01848 (2022).

12. Sali, R., Moradinasab, N., Guleria, S., Ehsan, L., Fernandes, P., Shah, T. U., Syed, S., & Brown, D. E. (2020). Deep Learning for Whole-Slide Tissue Histopathology Classification: A Comparative Study in the Identification of Dysplastic and Non-Dysplastic Barrett’s Esophagus. Journal of Personalized Medicine, 10(4), 141.

13. Graham, S., Vu, Q. D., Raza, S. E. A., Azam, A., Tsang, Y. W., Kwak, J. T., & Rajpoot, N. (2019). Hover-net: Simultaneous segmentation and classification of nuclei in multi-tissue histology images. Medical Image Analysis, 58,101563. 10.1016/j.media.2019.101563

14. Graham, Simon, et al. “Lizard: A Large-Scale Dataset for Colonic Nuclear Instance Segmentation and Classification.” Proceedings of the IEEE/CVF International Conference on Computer Vision

15. Mahfuz M, Das S, Mazumder RN, et al. Bangladesh Environmental Enteric Dysfunction (BEED) study: Protocol for a community-based intervention study to validate non-invasive biomarkers of environmental enteric dysfunction. BMJ Open. 2017;7(8). doi:10.1136/bmjopen-2017-017768

16. Human Cell Atlas. https://www.humancellatlas.org/learn-more/#event-launch-of-the-human-cell-atlas

17. C ZI. Mapping the Early Childhood Gut Across Ancestry , Geography and Environment. Published 2021. https://chanzuckerberg.com/science/programs-resources/single-cell-biology/pediatric-networks/mapping-the-early-childhood-gut-across-ancestry-geography-and-environment/

18. Zilbauer M, James KR, Kaur M, et al. A Roadmap for the Human Gut Cell Atlas. Nat Rev Gastroenterol Hepatol. 2023;20(9):597–614. doi:10.1038/s41575-023-00784-1

19. Syed S, Al-Boni M, Khan MN, et al. Assessment of Machine Learning Detection of Environmental Enteropathy and Celiac Disease in Children. JAMA Netw Open. 2019;2(6):e195822. doi:10.1001/jamanetworkopen.2019.5822

20. Khan M, Jamil Z, Ehsan L, et al. Quantitative Morphometry and Machine Learning Model to Explore Duodenal and Rectal Mucosal Tissue of Children with Environmental Enteric Dysfunction. Am J Trop Med Hyg. 2023;108(4):672–683. doi:10.4269/ajtmh.22-0063

21. Ewels P, Magnusson M, Lundin S, Käller M. MultiQC: Summarize analysis results for multiple tools and samples in a single report. Bioinformatics. 2016;32(19):3047–3048. doi:10.1093/bioinformatics/btw354

22. Dobin A, Davis CA, Schlesinger F, et al. STAR: Ultrafast universal RNA-seq aligner. Bioinformatics. 2013;29(1):15–21. doi:10.1093/bioinformatics/bts635

23. Love MI, Huber W, Anders S. Moderated estimation of fold change and dispersion for RNA-seq data with DESeq2. Genome Biol. 2014;15(12):1–21. doi:10.1186/s13059-014-0550-8

24. Leek JT. Svaseq: Removing batch effects and other unwanted noise from sequencing data. Nucleic Acids Res. 2014;42(21):e161. doi:10.1093/nar/gku864

25. Zhu A, Ibrahim JG, Love MI. Heavy-Tailed prior distributions for sequence count data: Removing the noise and preserving large differences. Bioinformatics. 2019;35(12):2084–2092. doi:10.1093/bioinformatics/bty895

26. Liberzon A. A description of the Molecular Signatures Database (MSigDB) Web site. Methods Mol Biol. 2014;1150:153–160. doi:10.1007/978-1-4939-0512-6_9

27. Liberzon A, Subramanian A, Pinchback R, Thorvaldsdóttir H, Tamayo P, Mesirov JP. Molecular signatures database (MSigDB) 3.0. Bioinformatics. 2011;27(12):1739–1740. doi:10.1093/bioinformatics/btr260

28. Liberzon A, Birger C, Thorvaldsdóttir H, Ghandi M, Mesirov JP, Tamayo P. The Molecular Signatures Database Hallmark Gene Set Collection. Cell Syst. 2015;1(6):417–425. doi:10.1016/j.cels.2015.12.004

29. Lundberg SM, Lee SI. A unified approach to interpreting model predictions. Advances in Neural Information Processing Systems. 2017;30.

